# Induction and characterisation of Aβ and tau pathology in *App^NL-F/NL-F^* mice following inoculation with Alzheimer’s disease brain homogenate

**DOI:** 10.1101/2024.07.11.602448

**Authors:** Silvia A. Purro, Michael Farmer, Emma Quarterman, Julia Ravey, David X. Thomas, Elizabeth Noble, Catherine Turnbull, Jacqueline Linehan, Tamsin Nazari, Sebastian Brandner, Mark A. Farrow, Dominic M. Walsh, John Collinge

## Abstract

Alzheimer’s disease (AD) is defined by the accumulation of neurofibrillary tangles containing hyperphosphorylated Tau and plaques containing Amyloid-β (Aβ). The aggregation of these two proteins is considered central to the disease. The lack of animal models that can recapitulate Aβ and tau pathologies without overexpressing these proteins has hindered AD research. Accelerating pathology by inoculating Aβ and tau seeds has helped to understand their prion-like propagation in the brain. Previous studies failed to characterise both Aβ and tau pathologies *in vivo* upon inoculating AD brain homogenates. Here we present a longitudinal and systematic study; we inoculated the *App^NL-F/NL-F^* knockin mice, which express humanised Aβ and murine wild-type tau, with extracts from diseased human brains to analyse the contribution of Aβ and tau assemblies to AD pathogenesis. We found that mice inoculated with AD brain extracts evinced early and prominent amyloid deposition, while those injected with control brain extracts or vehicle did not. Parenchymal and vascular amyloid accumulated in the same brain regions affected in control-inoculated *App^NL-F/NL-F^* mice. However, the extent of vascular amyloid far exceeded that seen in *App^NL-F/NL-F^*mice injected with control brain extracts, and parenchymal deposits extended to a previously untargeted brain region – the cerebellum. An end-point titration of an AD brain homogenate in *App^NL-F/NL-F^* mice demonstrated that human Aβ seeds can be titrated in a prion-like fashion, which is useful for sample comparison, diagnostic and risk studies. Notably, the inoculation of *App^NL-F/NL-F^* mice with AD brain homogenate induced intense tau phosphorylation, and provides more detailed context for the inoculation of *App^NL-F/NL-F^* mice with human samples to study temporal and mechanistic relationships between Aβ and tau pathology, vascular amyloid deposition and bioactivity of Aβ seeds.

## Introduction

Alzheimer’s disease (AD) is characterised by extracellular neuritic plaques, composed mainly of fibrillar amyloid-β (Aβ), and intracellular neurofibrillary tangles, which are insoluble deposits of hyperphosphorylated tau (Lane, Hardy et al. 2018). Aβ and tau exhibit specific prion-like properties, as they can propagate by templated seeding, spread within the brain, and from peripheral sites to the brain, induce neurotoxicity and resist inactivation to standard sterilisation techniques (Eisele, Bolmont et al. 2009). Recently, we presented evidence of iatrogenic AD in patients who received cadaveric-derived human growth hormone which contained Aβ seeds and tau (Purro, Farrow et al. 2018, Banerjee, Farmer et al. 2024). This human study highlighted the relevance of prion paradigms in neurodegenerative diseases, especially in AD. It is unclear if the administration of Aβ seeds alone is sufficient to induce tau pathology. Intracerebral inoculation of AD brain homogenate allows to study the onset and progression of Aβ and tau pathologies from a fixed point of time and a specific location in the brain, akin to classical prion transmission studies (Collinge 2016). Previous studies inoculating AD brain homogenate did not analyse tau pathology, or found little tau pathology when using models expressing wild-type tau (Ruiz-Riquelme, Lau et al. 2018, McAllister, Lacoursiere et al. 2020). Most of the Aβ transmission studies were performed using transgenic mice overexpressing proteins, which can cause unrelated disease events (Saito, Matsuba et al. 2014). Therefore, we examined the temporal and spatial dynamics of Aβ and tau deposits in the *App^NL-F/NL-F^* (NLF) mice, which express humanised Aβ and murine wild-type tau, upon inoculating AD brain extract containing Aβ and tau seeds. Here we present a longitudinal analysis that outlines the progression of both Aβ and tau deposition following the injection of AD brain extract to understand how these two prion-like proteins co-manifest their respective seeding abilities *in vivo*.

Given the elusive nature of Aβ seed structure (Ono and Watanabe-Nakayama 2021, Ulm, Borchelt et al. 2021), which currently precludes its specific biochemical quantification, we performed an end-point titration of an AD brain extract in the NLF mice. This method allowed us to quantify Aβ seeds in a tissue sample. We show that, similarly to prions, Aβ seeds can be titrated allowing different samples to be studied and compared for research, diagnoses, risk assessment and surveillance.

This study presents a systematic comparative analysis following intracerebral injection of human AD brain homogenate into NLF mice up to 480 days post-inoculation. We found significant differences of parenchymal Aβ load within the cerebellum, and blood vessels featuring vascular Aβ in mice inoculated with AD brain extract, compared to those inoculated with vehicle or human non-diseased brain homogenate. Quantifying vascular Aβ and cerebellar Aβ plaques allowed the titration of seeding activity (SD_50_) in AD brain homogenates with this model. Furthermore, our findings suggest a correlation between late tau aggregation and regions affected by early parenchymal Aβ in the AD-inoculated animals. Our work establishes NLF mice as a sensitive model to study vascular and parenchymal Aβ deposition *in vivo* and their relationship with tau pathology, supporting their role as a pathophysiologically relevant bioassay tool in the study of AD.

## Materials and methods

### Use of human tissues and research ethics

This study was performed with Ethics approval from the North East - Newcastle & North Tyneside 2 Research Ethics Committee; REC reference: 11/NE/0348 and London Queen Square Research Ethics Committee REC reference: 03/N038. Storage and biochemical analysis of post-mortem human tissue samples and transmission studies to mice were performed with written informed consent from patients with the capacity to give consent or a relative in accordance with applicable UK legislation and Regulatory Codes of Practice. One control case with no signs of neurodegenerative disease and three pathologically Alzheimer’s disease diagnosed cases were kindly provided under a material transfer agreement from the Oxford Brain Bank, Oxford University Hospitals NHS Trust and the Queen Square Brain Bank for Neurological Disorders, UCL Institute of Neurology (Table 1).

**Table 1.** Demographic details of the cases.

| **Patient** | **Healthy Control** | **AD 1** | **AD 2** | **AD 3** |
| --- | --- | --- | --- | --- |
| Age of onset | N/A | 64 | 65 | 57 |
| Age of death | 41 | 77 | 76 | 64 |
| Disease duration (years) | N/A | 13 | 11 | 7 |
| Gender | Male | Male | Female | Male |
| Brain region | Frontal cortex | Frontal cortex | Frontal cortex | Frontal cortex |
| Neuropathological diagnosis | “Healthy” control – no evidence of neurodegenerative disease. Acute hypoxic ischaemic Damage consistent with the history of a relatively fast cardiac death. | AD long history. Braak VI. Severe CAA Lewy bodies TDP43 proteinopathy. | AD long history. Braak VI. Severe CAA TDP43 proteinopathy. | AD long history. Braak VI. Moderate CAA. |

Frontal cortex grey matter from brain tissue was homogenised with glass grinders in Dulbecco’s phosphate buffered saline lacking Ca++ or Mg++ ions (PBS) at 10 % (w/v) and subsequently diluted to 1 % (w/v) in PBS to inoculate the mice.

### Multiplex Aβ peptide immunoassay

Levels of Aβ peptides in the human brain homogenates were determined using the V-Plex Aβ peptide panel (6E10) immunoassay from Meso Scale Discovery (MSD) according to the manufacturer’s instructions. All standards and samples were diluted in PBS. Total fraction (1 % (w/v) brain homogenate) was spun at 13,000 rpm for 15 min at 4 °C and the supernatant labelled as soluble fraction. Before analysis, total and soluble fractions of the homogenate were incubated with guanidinium-HCl (GuHCl) (6 M final concentration) to disaggregate preformed Aβ aggregates. Aβ peptide levels were determined reading the plate immediately using a QuickPlex SQ 120 and data analysed using MSD workbench software.

### Preparation of synthetic peptides used for Western blotting (WB)

Lyophilised Aβ_1-42_ and Aβ_1-40_ were dissolved in 50 mM Tris-HCl, pH 8.5, containing 7 M GuHCl and 5 mM EDTA at a concentration of 1 mg/ml and incubated at RT overnight. Samples were then centrifuged for 30 min at 16,000 g and chromatographed on a Superdex 75 10/300 column eluted at 0.5 ml/min with 50 mM ammonium bicarbonate, pH 8.5. The concentration of the peak fraction for each sample was determined by absorbance at 275 nm. The peptide was then diluted to 10 ng/μl, and stored frozen at −80 °C in aliquots.

### Western blotting (WB) of Aβ in brain extracts

Crude 1% (w/v) brain homogenate in PBS was boiled in 2X sample buffer (100 mM Tris, 4 % w/v SDS, 24 % v/v glycerol with 0.02 % (w/v) phenol red) for 10 min at 100 °C, then electrophoresed on hand-poured, 15-well 16 % polyacrylamide Tris-tricine gels. Aβ_1-42_ and Aβ_1-40_ were run as loading controls and protein transferred onto 0.2 μm nitrocellulose at 400 mA and 4 °C for 2 hr. Blots were microwaved in PBS and Aβ detected using the anti-Aβ antibodies 3D6, 6E10, m266, 4G8, HJ2 and HJ7.4, and bands visualized using ECL Plus substrate and X-ray film.

### Mouse transmission studies

Work with mice was performed under approval and licence granted by the UK Home Office (Animals (Scientific Procedures) Act 1986); Project Licence 70/9022 and conformed to University College London institutional and ARRIVE guidelines (http://www.nc3rs.org.uk/ARRIVE/).

NLF knock-in mice (*App^NL-F/NL-F^*) containing the Aβ domain humanised and carrying the Swedish and Beyreuther/Iberian mutations were maintained on a C57BL/6J background and used as homozygotes. Mouse genotype was determined by PCR of ear punch DNA and mice were uniquely identified by sub-cutaneous transponders. Mice at 6-8 weeks were assigned to experimental groups, anaesthetised with halothane and O_2_ and inoculated intracerebrally into the right lobe with 30 µl of a 1 % (w/v) human brain homogenate prepared in PBS or vehicle (PBS) alone. At stated times, mice were anesthetized with isoflurane/O_2_, decapitated and brains were collected. The left hemisphere was fixed for immunohistochemistry and the right hemisphere was frozen for biochemistry. A subset of 5 brains per group from 4 to 12 months post inoculation were collected whole, fixed and coronal cuts at four different levels were examined histopathologically.

### Immunohistochemistry (IHC)

The left hemisphere was fixed in 10 % (w/v) buffered formal saline followed by incubation in 98 % (w/v) formic acid for 1 hour. After washing, brains were processed and paraffin wax embedded. Serial sections of 5 μm nominal thickness were taken. Aβ deposition and tau pathology (phosphorylated-Tau) were visualized using biotinylated 82E1 (cat n.10326, IBL, Japan) and AT8 (pTau) (#MN1020B, Invitrogen), respectively, as the primary antibody, using a Ventana Discovery automated immunohistochemical staining machine (Roche, Burgess Hill, UK) and proprietary solutions. Visualization was accomplished with development of 3’3 diaminobenzidine tetrahydrochloride as the chromogen (DAB Map Kit, Ventana Medical Systems). Haematoxylin was used as the counterstain.

### Quantification of AT8 deposits, vascular and parenchymal Aβ immunoreactivity

Histological slides were digitised either on a LEICA SCN400F scanner (LEICA Milton Keynes, UK) as described previously (Jaunmuktane, Mead et al. 2015) or on a Hamamatsu Nanozoomer S360.

Digital image analysis on whole slides was performed using Definiens Developer XD 2.6 (Definiens, Munich, Germany). Vascular and parenchymal Aβ were quantified as before (Purro, Farrow et al. 2018), quantifications were performed blind to experimental group. Briefly, either one paramedian (approximately 200 µm) sagittal section or four coronal sections per mouse were analysed. For vascular Aβ, blood vessels in six anatomical areas of the meninges covering the dorsal part of the brain, including the olfactory bulb and the cerebellum, were analysed by manually counting the number of Aβ negative and positive vessels. For parenchymal Aβ quantification, brightness thresholds were used to identify the tissue within each image and relevant brain regions were selected as distinct regions of interest (ROI) to separate cortex, hippocampus, and cerebellum; larger artefacts were also manually selected for exclusion from analysis. Brown and blue stain levels were then calculated and combined to give 3 stain representations: brown staining, blue staining and brown^+ve^ staining. All areas with brown stain intensity greater than 0.15 arbitrary units (au) and brown^+ve^ greater than 0.1 as were subcategorised to light brown < 0.5 au <= dark brown. Dark brown areas were then grown into light brown areas to give potential plaques which were then accepted or excluded based on size, shape and composition criteria.

QuPath software was used to analyse AT8 specific staining. Each image was set to Brightfield_H_DAB. To outline each brain area, a pixel classification thresholder was created, with the following parameters; resolution = moderate, Channel = Hematoxylin, Prefilter = Gaussian, smoothing sigma = 1.7, threshold = 0.08, above threshold = region, below threshold = unclassified, region = everywhere. To create annotations from the pixel classier, the minimum object size = 500,000 µm^2^, minimum hole size = 400 µm^2^, where objects were split and objects were created for ignored classes. To detect AT8 stain, an object classifier was created. Each annotation from the previous script was selected for each image. A positive cell detection was ran with the following parameters; pixel size = 0.2 µ, background radius = 10 µ, sigma = 1.0 µ, minimum area = 2.0 µ, maximum area = 100 µ, threshold = 4.0 and maximum background = 2.0. Then from each positive cell detection, an object classier was trained to detect positive cell detections stained with AT8, as well as which detections were negative. Artefacts were removed based on colour, shape and size.

### NLF brain homogenisation and fractionation

Weighed hemibrains were homogenised in PBS using a Precellys 24 ribolyser (Bertin Technologies) (6,500 rpm, 1 cycle, 45 sec) to prepare a 20 % (w/v) homogenate, using zirconia oxide ceramic beads (#15515809). The homogenate was diluted 1:1 with PBS with 2 x Protease/phosphatase inhibitors (2 tablets Thermo Scientific Pierce # A32961 per 10ml PBS). A portion of the resulting 10 % (w/v) homogenate was further fractionated by centrifugation at 1000 x g and 4° C for 5 min. The supernatant which referred to as “the total fraction” was removed and stored at −80 °C, or further fractionated by centrifugation at 75,000 rpm and 4 °C for 30 min using a TLA-110 rotor in Optima Max-XP centrifuge. The supernatant, referred to as “the soluble fraction” was removed and stored at −80 °C.

### Aβ_x-42_ MSD assays (V-PLEX Human Aβ42 Kit # K151LBG)

Samples were incubated with GuHCl (final concentration of 6 M) for 15 min at RT and then diluted as appropriate in PBS. An eight-point standard curve was prepared from the Aβ_1-42_ peptide in PBS. The microplate was blocked for 1 hr at RT (600 rpm) and then washed 3 times with PBS (0.05 % (w/v) Tween). Samples and standards were added in duplicate to the microplate 50 µl/well and incubated at RT (600 rpm) for 1hr. The plate was washed 3 times with PBS (0.05 % (w/v) Tween) and detection antibody was added 25 µl/well (diluted 1:50). The antibody was incubated at RT (600 rpm) for 1 hr and then washed 3 times with PBS (0.05 % (w/v) Tween). 150 µl/well 2X read buffer added. The plate read immediately using a QuickPlex SQ 120 and data analysed using MSD workbench software.

### Endpoint titration assay to estimate SD_50_

NLF knock-in mice (*App^NL-F/NL-F^*) at 6-8 weeks were assigned to experimental groups, anaesthetised with halothane and O_2_ and inoculated intracerebrally into the right lobe with 30 µl of AD2 brain homogenate serially diluted (1% to 0.001% w/v) in control human brain homogenate, or 30 μl of 1% w/v unaffected control human brain homogenate (n = 5 mice per group). After 120 days post inoculation, mice were anesthetized with isoflurane/O_2_, decapitated and brains were collected and fixed for immunohistochemistry. One sagittal slide per mouse was cut and stained with 82e1-biotinylated antibody to quantify parenchymal and vascular Aβ as mentioned above. Animals scoring >0 % of plaques in the cerebellar region where classified as positive seeded. Independently, an animal was rated positive if presented one or more dorsal blood vessels with Aβ depositions (vascular Aβ). The experimenter performed both quantifications blinded to the treatments. We calculated the Difference of logarithms or proportional distance = [(% of induced animals at the last dilution where induction rate is higher than 50%)-50%]/[(% of induced animals at the last dilution where induction rate is higher than 50%)-(% of induced animals at the last dilution where induction rate is lower than 50%)] and then: log_10_ 50% end point dilution = log_10_ of the last dilution where induction rate is higher 50% - (difference of logarithms × logarithm of dilution factor) (Reed and Muench 1938). This method was applied for vascular and parenchymal Aβ deposits in the cerebellar region, independently.

### Statistical Analysis

All statistical analysis and graphs were generated using the package GraphPad PRISM v6 (GraphPad Software, Inc., La Jolla, USA). Error bars on graphs denote standard deviation, with statistical significance determined by One-way ANOVA followed by Dunnett’s multiple comparison test (two-tailed) unless specified differently. Statistical significance was set to P < 0.05.

### Data availability

Data will be available upon reasonable request. Correspondence and requests for materials should be addressed to J.C.

## Results

### NLF mice inoculated with homogenates of AD brain exhibit accelerated accumulation of parenchymal and vascular amyloid

Many AD mouse models present artefacts due to overexpression of APP and/or the use of an exogenous promoter (Saito, Matsuba et al. 2014) therefore, we decided to perform our transmission experiments using the NLF knockin mice (Saito, Matsuba et al. 2014). We inoculated female mice via intracerebral injection at 6 to 8 weeks old with 30 µl of human brain homogenate control, AD1, AD2, AD3 (Table 1) or vehicle (PBS). Mice were culled at the following time points: 2, 7, 15, 30, 45, 60, 90, 120, 240, 360, 480 days post-inoculation (dpi). For short time points (2 to 90 dpi) groups were of 5 mice and for longer time points (from 120 dpi onwards) groups contained 15 animals to mitigate for intercurrent diseases. Brain hemispheres were collected either for biochemistry or histology in order to carry out a systematic analysis of plaques, amyloid deposition in blood vessels and quantification of Aβ peptides.

Vehicle- and control brain homogenate-inoculated mice were indistinguishable. Control- and PBS-inoculated mice had no discernible Aβ immunoreactivity (IR) until 120 dpi and at this time amyloid was only detectable in the cerebral cortex area with all other regions free of Aβ (Fig. 1, Suppl Fig 1). However, after 240 dpi Aβ IR was evident in the cerebral cortex and hippocampus of all control- and PBS-inoculated mice. Occasional Aβ IR was detected in the olfactory bulb and tectum, but not in other brain regions. Thereafter, amyloid progressively covered the hippocampus, corpus callosum, cerebral cortex, and then olfactory bulb. At 360 dpi, some ventral regions were affected, along with a few blood vessels on the dorsal meninges (Fig. 1, Suppl Fig 1).

**Figure 1.**
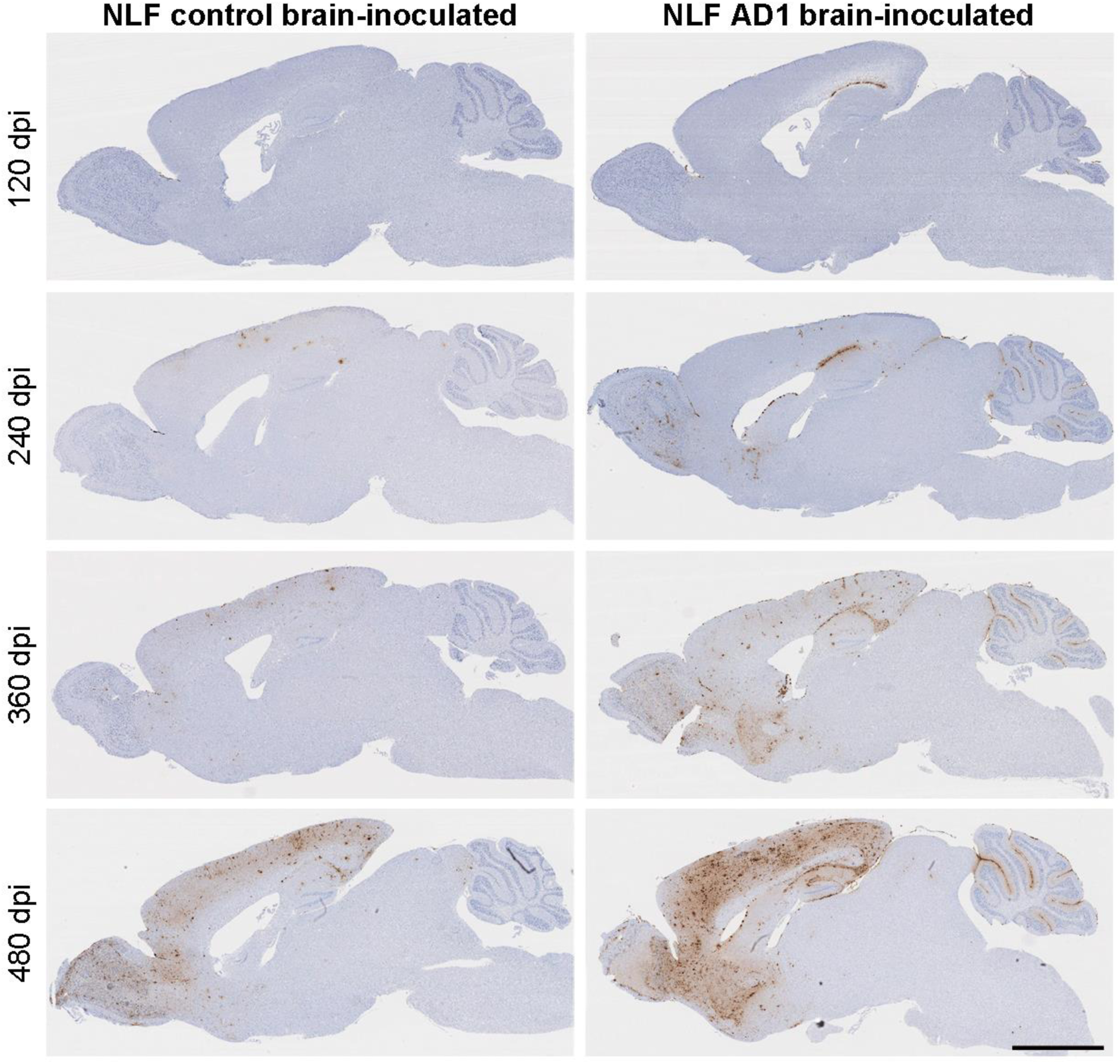
Inoculation of NLF mice with human AD brain homogenate accelerates amyloid accumulation. Mice were intracerebrally inoculated with either control or AD brain and culled at the times indicated. Aβ deposition was assessed on sagittal sections stained using an anti-Aβ antibody, a biotinylated derivative of the 82E1. Scale bar = 2 mm. dpi = days post-inoculation.

AD1-inoculated mice culled at 2-15 dpi presented no traces of the inoculum and there were no amyloid deposits at this stage, demonstrating that there is no detectable persistence of the inoculum in the tissue (Suppl. Fig 2). At the 30, 45 and 60 dpi AD-inoculated mice (n=5 mice per time point) began to show small puncta of Aβ IR in the corpus callosum (Suppl. Fig. 2). Some of these samples (4 out of 15) had amyloid depositions in one or two dorsal meningeal blood vessels surrounding the cerebral cortex. At 30 dpi, 2 out of 5 AD1-inoculated mice had occasional parenchymal IR and/or a couple of Aβ positive blood vessels, but by 60 dpi all the animals (5 out of 5) had parenchymal and/or vascular IR. From 120 dpi on, AD1-inoculated mice had Aβ IR in the hippocampus, cerebral cortex and cerebellum. Later on, at 240, 360 and 480 dpi, parenchymal amyloid progressively covered all areas starting from the corpus callosum, cerebellum, cerebral cortex and hippocampus, with later involvement of the olfactory bulb and ventral areas of the brain (anterior olfactory bulb, striatum, hypothalamus, medulla, anterior commissure). Similarly, more blood vessels gradually had amyloid deposits beginning with cortical meningeal vessels on the cerebral cortex, then the vessels in the cerebellar sulci and vessels near the hippocampus. Later this extended to include meningeal vessels near the olfactory bulb from dorsal to ventral regions (Fig. 1).

We collected brain samples at 120, 240 and 360 dpi and performed coronal cuts at three levels (approximate Bregma coordinates −6.2, −1.7 and 1.1 mm), and cut four histological sections from each mouse brain. One slice from each section was stained to quantify Aβ IR in each hemisphere. Amyloid load on the ipsilateral side was similar to the contralateral hemisphere in all time points regardless of the inoculum used (Suppl Figure 3).

### Inoculation with homogenates from two other AD brains accelerates amyloidosis in NLF mice measured by both IHC and MSD-based immunoassay

To test the generalisability of the induction seen with AD1 brain, we used two methods, IHC and MSD-based immunoassay.

We compared amyloid deposition in brains of mice inoculated with homogenates of AD1, AD2 and AD3 versus control and PBS vehicle (Table 1). At each of the time points, we quantified parenchymal amyloid in whole sagittal section. Dorsal vascular amyloid was common in the brains of all NLF mice inoculated with AD-brain extracts after 120 dpi. In the cerebellum, Aβ depositions most frequently occurred in the outermost molecular layer. All AD-inoculated groups had a significantly higher plaque load than PBS-inoculated mice at equivalent time points up to 480 dpi (Fig. 2). At 120, 240 and 360 dpi PBS- and control-inoculated mice had only 10% of the total area covered by amyloid whereas in brains from AD-inoculated mice amyloid burden was at least 10-fold greater (area covered by plaques at 120 dpi: average controls (PBS+C) = 0.01 % vs AD1-3 = 0.1 %; at 240 dpi: average controls = 0.14 % vs AD1- 3 = 1.5 %; at 360 dpi: average controls = 0.5 % vs AD1-3 = 0.52 %). At 480 dpi, this difference was less noticeable, with AD-inoculated mice having approximately 2-fold more Aβ than mice inoculated with PBS or controls brain homogenate (at 480 dpi: average controls = 8.5 % vs AD1-3 = 22.6 %). Similar trends were seen when corresponding hemi brains were analysed using the MSD-based Aβ_42_ immunoassay. All groups inoculated with AD brain extracts had significantly more Aβ_x-42_ than control mice at the time points tested (Fig. 3).

**Figure 2.**
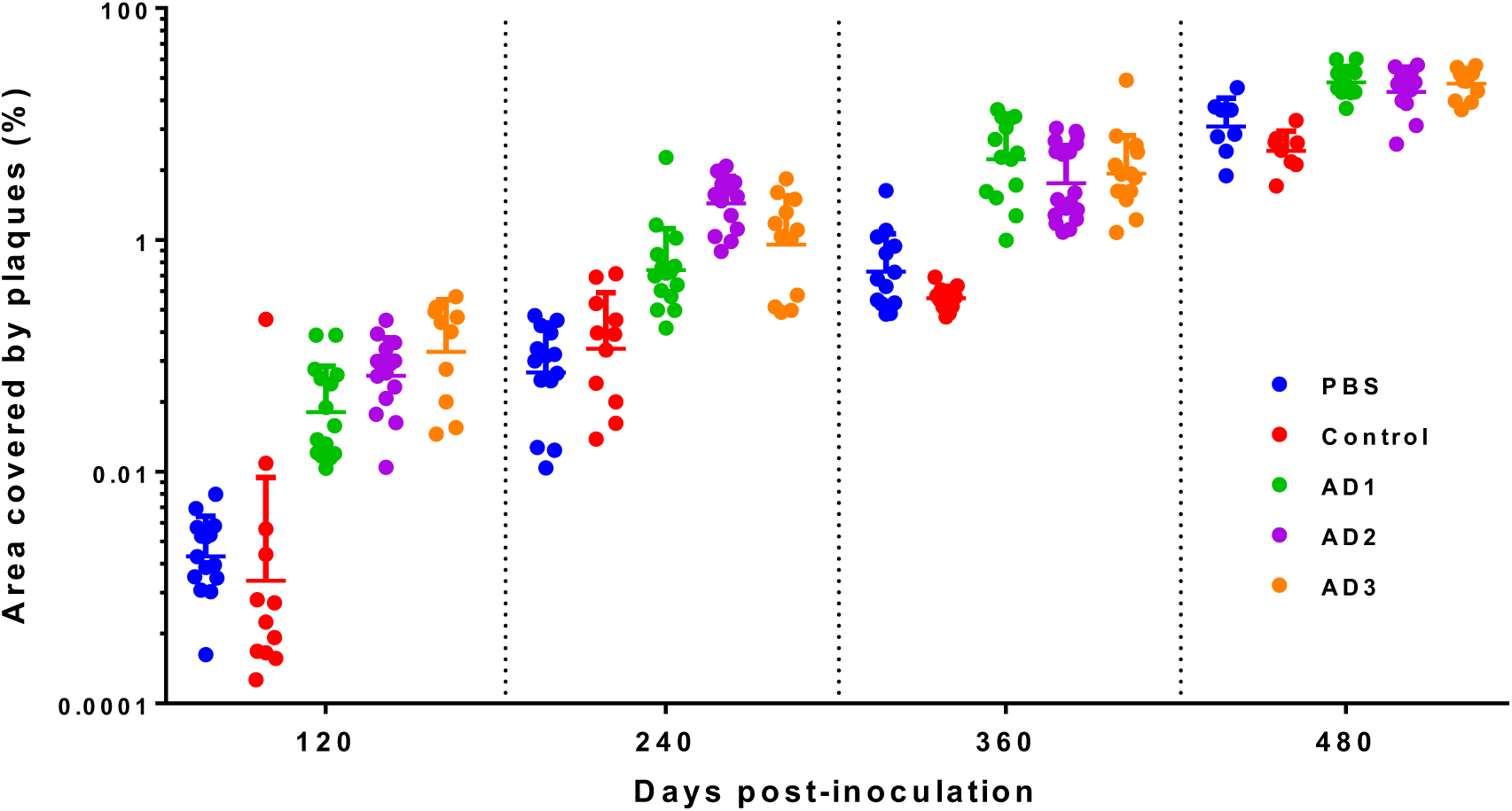
Inocula produced from three different AD brains similarly accelerate accumulation of Aβ. NLF mice were intracerebrally inoculated with either control or AD brain homogenate, and then culled at stated times post inoculation. Sagittal brain sections were stained using biotinylated 82E1 and immunoreactivity quantified. Highly significant (adjusted P<0.0001) differences were found between controls and ADs inoculated samples at 120, 240 and 360 days post-inoculation. At 480 days post-inoculation differences where smaller but still statistically significant (PBS vs AD1 P=0.0003, PBS vs AD2 P=0.004 PBS vs AD3 P=0.0003, One way ANOVA followed by Dunnett’s multiple comparison test). There was no significant difference between PBS and control brain inoculated mice at equivalent time points. Statistics were carried out on log-transformed data. 120 dpi n=10-15 mice, 240 dpi n=11-15 mice, 360 dpi n=11-19 mice, 480 dpi n=8-11 mice per inoculum. Error bars = Mean ± SD.

**Figure 3.**
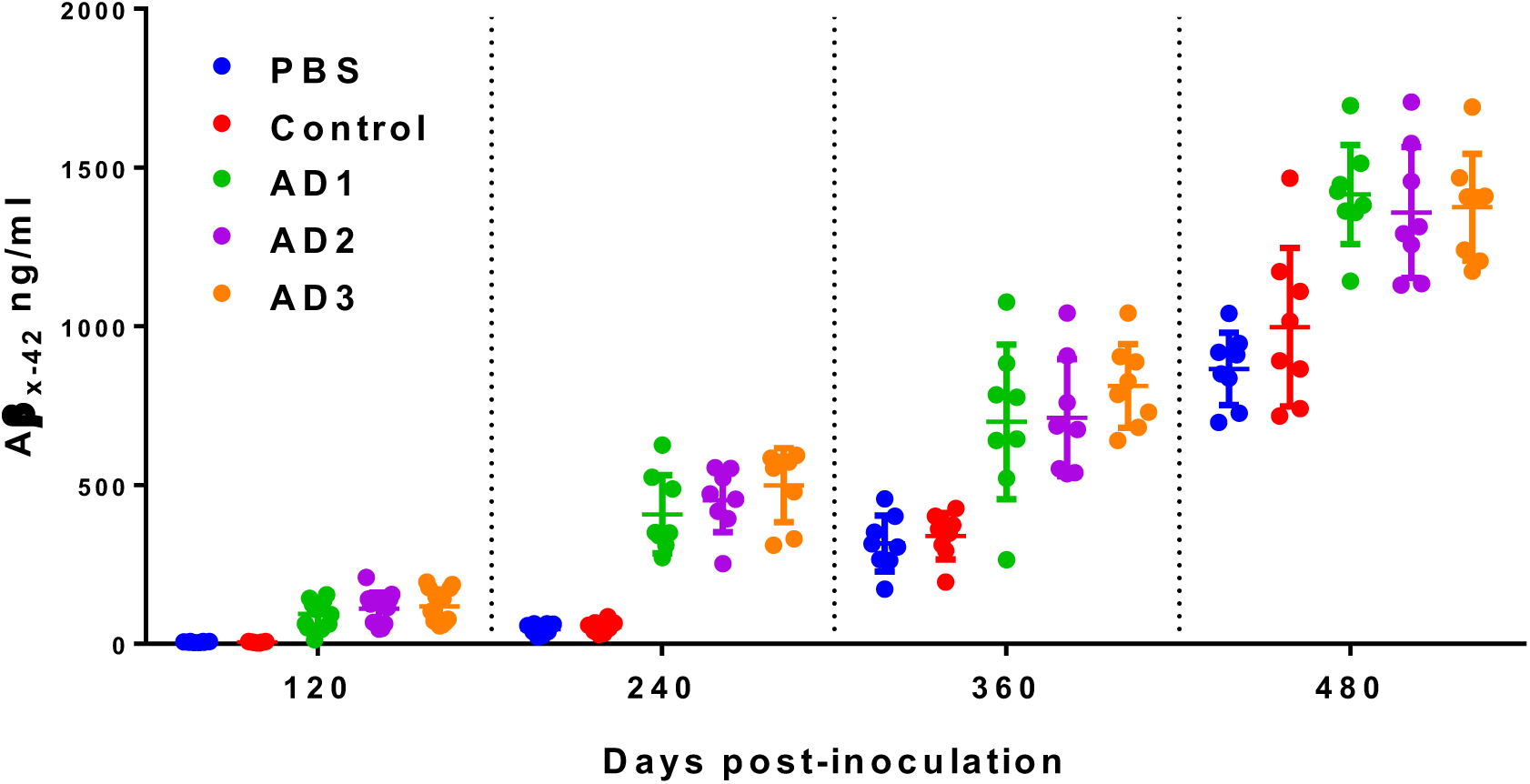
Quantification of Aβ_42_ levels in total mouse brain homogenate by MSD assay. One way ANOVA followed by Dunnett’s multiple comparison test shows highly significant (adjusted P<0.0001) differences between all groups inoculated with AD brain homogenate and PBS inoculated mice for each time point. There was no significant difference between mice inoculated with control brain or PBS. Bars represent the mean concentration ± SD. N=8.

### AD brain extracts used to inoculate NL-F mice contain comparable levels of immunoassay measured Aβ

To investigate the forms of Aβ in the 3 AD brains used to inoculate NLF mice, we analysed homogenates using Western blotting and MSD-based immunoassays combined with centrifugation. For Western blotting we employed six anti-Aβ antibodies with epitopes ranging across the entire Aβ_1-42_ sequence (Suppl Figure 4A). We found that AD2 contained higher levels of Aβ measured by 3D6, 6E10, m266 and 4G8, but that AD1, AD2 and AD3 had comparable levels of Aβ_42_ (Suppl Figure 4B-C). Analysis of samples using MSD-based immunoassays before and after centrifugation, similarly indicated that AD2 contained the highest level of Aβ_40_ (Table 2), whereas AD1, AD2 and AD3 had comparable levels of Aβ_42_ (Table 2). Fractionation by centrifugation revealed that the majority of both Aβ_40_ and Aβ_42_ in the total homogenate were readily sedimented. Specifically, centrifugation at 1000 x g removed >90% of Aβ_40_ and Aβ_42_, and higher speed (16,000 x g) removed >99% of the MSD measured Aβ. These results indicate that only a tiny portion of Aβ in the inoculum injected into mice is readily diffusible. In future studies, it will be important to analyse fractions of AD brain homogenates and conduct titration experiments to relate seeding activity to the biochemical characteristics of Aβ in inocula.

**Table 2.** Quantification of Aβ peptides in brain homogenates used for inoculations Levels of Aβx-40 and Aβx-42 in total and soluble fractions of control and Alzheimer’s disease (AD1, AD2, AD3) human brain homogenates and APPPS1 mouse brain homogenate were quantified by immunoassay.

| Sample | Aβ 40<br>(ng/ml) | Aβ 42<br>(ng/ml) |
| --- | --- | --- |
| **10 % Brain homogenate, Total fraction** |  |  |
| Control | 7.44 | 1.26 ± 1.14 |
| AD1 | 274.26 ± 13.76 | 687 ± 12.21 |
| AD2 | Over limit | 644 ± 14.71 |
| AD3 | 200.36 ± 16.53 | 858 ± 8.23 |
| **10 % Brain homogenate, Clarified 1000 x g for 5 min** |  |  |
| Control | 0 ± 2.83 | 0 ± 0.68 |
| AD1 | 25.57 ± 1.36 | 39.1 ± 1.95 |
| AD2 | 122.4 ± 10.56 | 63.87 ± 6.42 |
| AD3 | 22.33 ± 2.10 | 68.53 ± 9.40 |
| **1 % Brain homogenate, Total fraction** |  |  |
| Control | 2.05 ± 0.64 | 6.67 ± 0.97 |
| AD1 | 13.67 ± 0.65 | 30.47 ± 2.40 |
| AD2 | 160.1 ± 10.75 | 43.33 ± 0.80 |
| AD3 | 14.20 ± 0.30 | 57.73 ± 2.33 |
| **1 % Brain homogenate, Soluble fraction. Supernatant 16000 x g for 15 min** |  |  |
| Control | 0.66 ± 0.25 | 0.27 ± 0.09 |
| AD1 | 1.34 ± 0.49 | 1.6 ± 0.22 |
| AD2 | 3.56 ± 0.17 | 1.14 ± 0.25 |
| AD3 | 1.82 ± 0.38 | 0.34 ± 0.05 |

### Inoculation of NLF mice with AD brain homogenate induces amyloid deposition in the cerebellum

Analysis of individual brain regions revealed Aβ IR in the cerebellum only of mice inoculated with AD brain extract (Fig. 4); animals inoculated with either PBS or human control brain homogenate had almost no detectable Aβ IR in the cerebellum even at the 480 dpi when Aβ IR was extensive in other areas of the brain. For example, control-inoculated samples displayed an average of 20.5 % amyloid coverage in hippocampus and cortex and 0.2 % in the cerebellar region compared to 46 % in hippocampus and cortex and 10.5 % in cerebellum for AD3-inoculated samples at 480 dpi. These results demonstrate that in addition to accelerating the deposition of amyloid in regions prone to aggregation of endogenous NLF Aβ, inoculating with homogenates from AD brain induces aggregation of Aβ in brain regions unaffected in uninoculated NLF mice.

**Figure 4.**
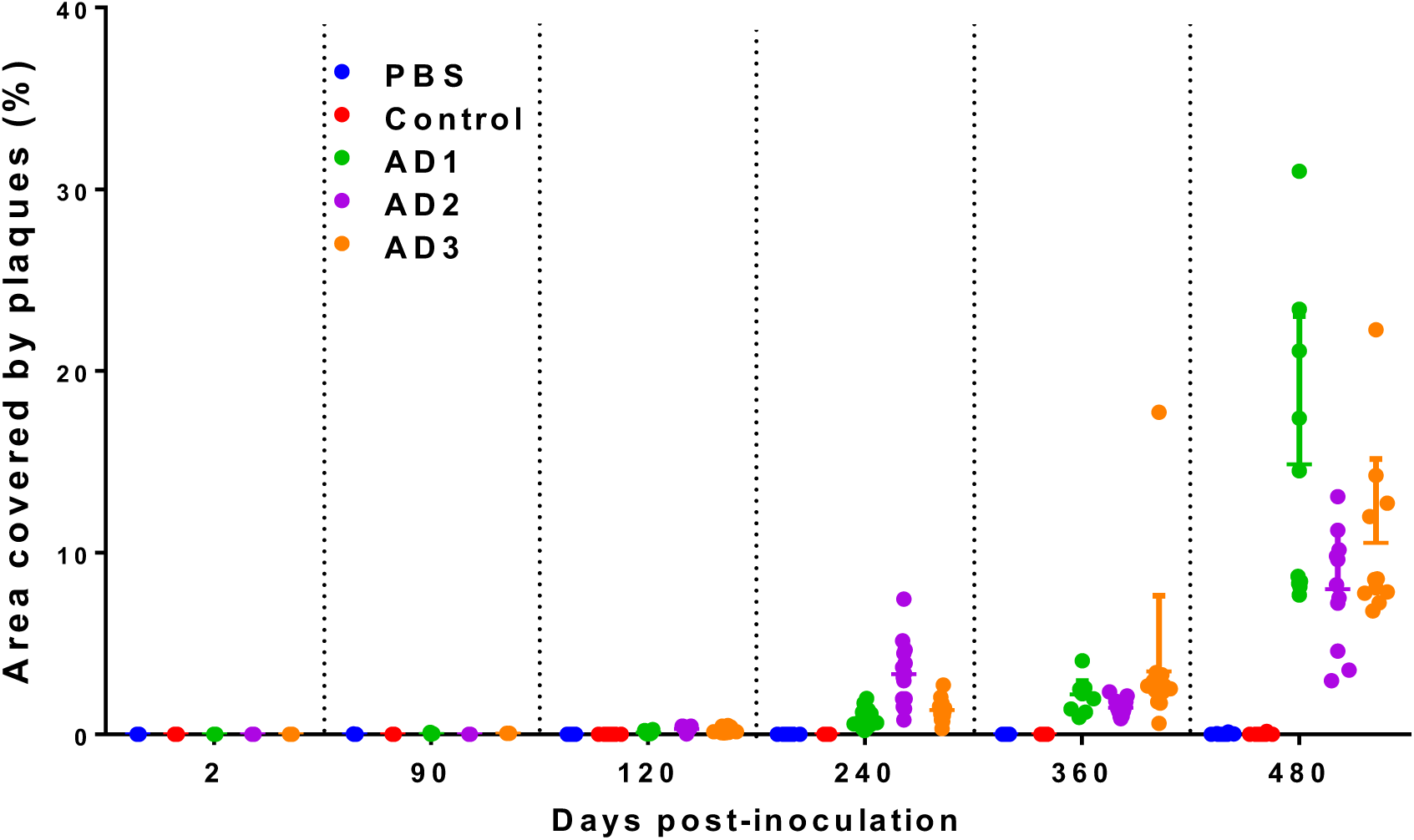
Inoculation of NLF mice with AD brain homogenate induces deposition of amyloid in the cerebellum. Graph shows the area covered by plaques (mean ± SD) expressed as a percentage of the total area following inoculation with homogenates of control or Alzheimer’s disease (AD1, AD2, AD3) brain and PBS. Sagittal sections were stained with biotinylated 82E1. Significant differences were found between PBS-inoculated and AD-inoculated mice at 90 dpi (PBS vs AD3 P=0.04), at 120 dpi (PBS vs AD1 P=0.001, PBS vs AD2-AD3 P<0.0001), at 240 dpi (PBS vs AD1 P=0.002, PBS vs AD2 P<0.0001, PBS vs AD3 P=0.0001), at 360 dpi (PBS vs AD1-AD3 P<0.0001, PBS vs AD2 P=0.006) and at 480 dpi (PBS vs AD1 P<0.0001, PBS vs AD2 P=0.01, PBS vs AD3 P=0.0015, Kruskal-Wallis followed by Dunn’s multiple comparison tests). There was no significant difference between PBS-and control brain-inoculated mice. 120 dpi n=10-15 mice, 240 dpi n=11-15 mice, 360 dpi n=11-19 mice, 480 dpi n=8-11 mice per inoculum.

### Inoculation of NL-F mice with AD brain accelerates the appearance and extent of vascular amyloid

Previously, we found that intracerebral inoculation of NLF mice with Aβ seeds can induce accumulation of amyloid in cerebral blood vessels (Purro, Farrow et al. 2018). Here, we systematically examined the rate of appearance and distribution of vascular amyloid from mice inoculated with AD versus control brain homogenates. From 90 dpi onwards all AD inoculated mice evinced Aβ in blood vessels, whereas vascular amyloid was only detected in the PBS and controls groups at 240 dpi (Fig. 5). These results confirm our previous findings that upon injection of Aβ seeds the blood vessels appear to be one of the first targets for amyloid deposition.

**Figure 5.**
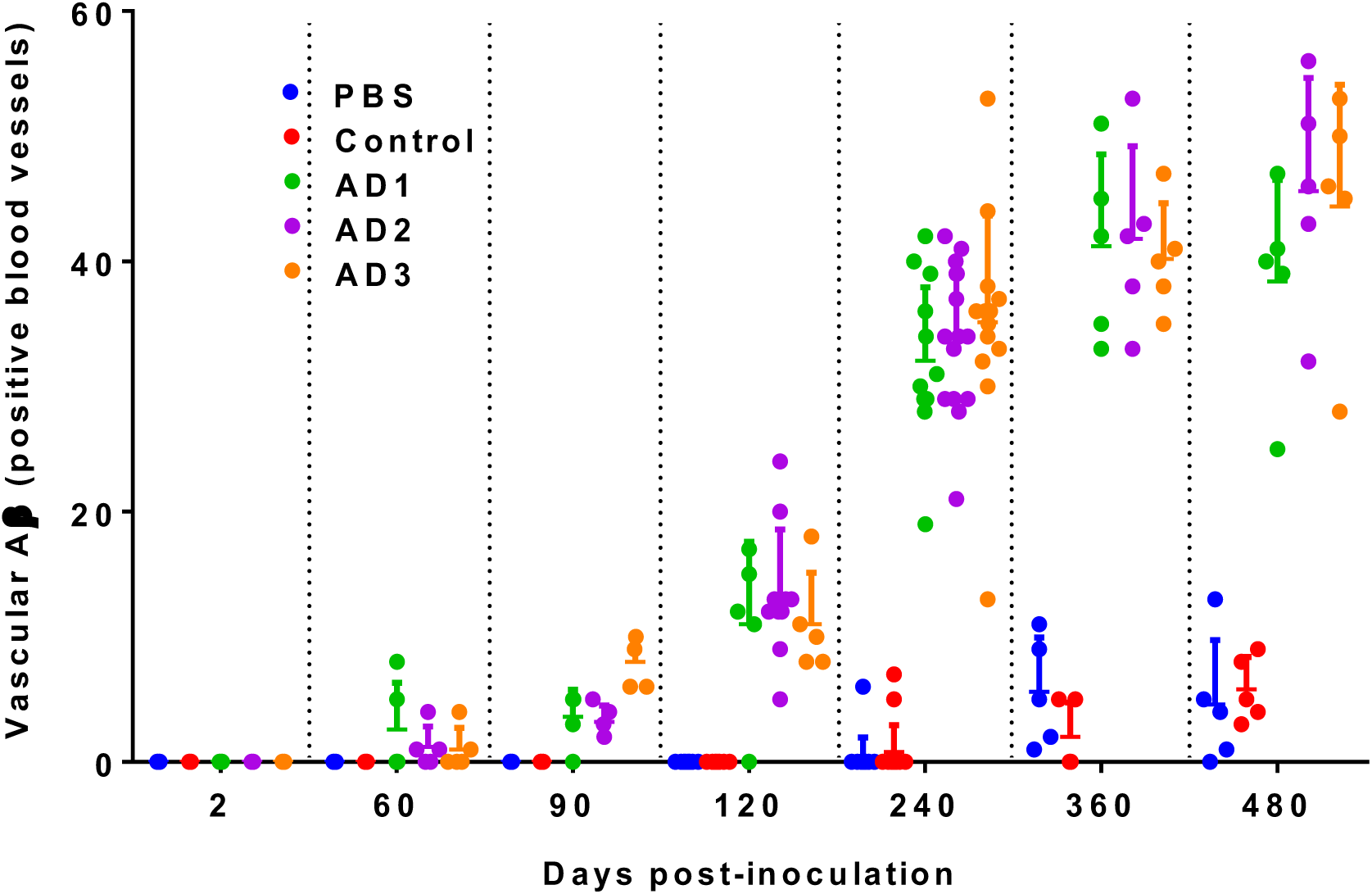
Inoculation of NLF mice with AD brain homogenate accelerates the appearance and extent of amyloid immunoreactivity in meningeal blood vessels. Sagittal sections were stained with biotinylated 82E1 and dorsal vessels with Aβ immunoreactivity counted. PBS-inoculated mice presented highly significant (adjusted P<0.0001) differences compared to mice inoculated with AD brain homogenates at each time post inoculation above 90 dpi (One way ANOVA followed by Dunnett’s multiple comparison test). At 90 dpi, PBS-inoculated mice were significantly different from (PBS vs AD1 P=0.002, PBS vs AD2 P<0.0001, PBS vs AD3 P=0.007). For those time points where all controls were scored zero, statistical significance was confirmed by the Kruskal-Wallis test. There were no significant differences between PBS- and control brain -inoculated mice. Mean ± SD is shown. 2 dpi n=5, 60 dpi n=5, 90 dpi n=5, 120 dpi n=5-10, 240 dpi n=14-15 mice, 360 dpi n=5, 480 dpi n=5 mice per inoculum.

To further explore the relationship between parenchyma and vascular amyloid, we plotted the number of Aβ positive blood vessels versus total plaque load or versus cerebellar plaque load per mouse inoculated (Suppl Fig 5). Vascular Aβ levels were weakly or not correlated with either. These results suggest that the deposition of Aβ in blood vessels does not follow the same pattern as the Aβ amyloid deposition in the parenchyma; these Aβ pathologies evolve separately. These results raise the possibility that other factors like Aβ conformation or assembly state (“strain type”), cell vulnerability and the differential diffusion of Aβ may contribute to the distinct evolution of amyloid at these sites (Qiang, Yau et al. 2017).

### End-point titration of Aβ seeds from AD brain homogenate

Studies quantifying the Aβ seeding activity present in biofluids and different human tissues would be useful for iatrogenic risk assessment and as a biomarker to measure the effect of new strategies against AD. As a proof of principle, we performed an end-point titration of AD2 brain homogenate in the NLF mice. Six to eight week old female NLF mice (n=5 mice per group) received one injection in the right parietal lobe of 30 μl of AD2 brain homogenate serially diluted (1% to 0.001% w/v) in control human brain homogenate, or 30 μl of 1% w/v control brain homogenate. Four months after injection, dorsal meningeal blood vessels with amyloid deposition (vascular Aβ count), total and cerebellar amyloid plaques were quantified for each inoculum (Figure 6). The highest levels of amyloid IR were seen with the 1 % (w/v) homogenate and the lowest with the 0.001 % (w/v) homogenate. The amount of both cerebellar parenchymal and vascular meningeal amyloid was strongly dependent on the dilution of the AD2 brain homogenate (Figure 6). The number of dorsal blood vessels that were Aβ IR positive was significantly different from control-inoculated mice when AD2 homogenate was diluted to 0.1 % (w/v) (p=0.01) (Figure 6). To quantitatively determine the AD2 brain homogenate seeding activity, we calculated the seeding dilution (SD_50_) at which half of the inoculated animals showed either vascular Aβ or Aβ depositions in the cerebellum using the Reed-Muench method (Reed and Muench 1938). The titers were similar when calculated using the data either from cerebellar and vascular Aβ amyloid: SD_50=_ 10^6.02^ and 10^6.35^ per gram of wet brain, respectively. These results demonstrate that relatively low levels of seeds are sufficient to induce amyloid formation in NLF mice and that this model is useful to bioassay Aβ seeding activity.

**Figure 6.**
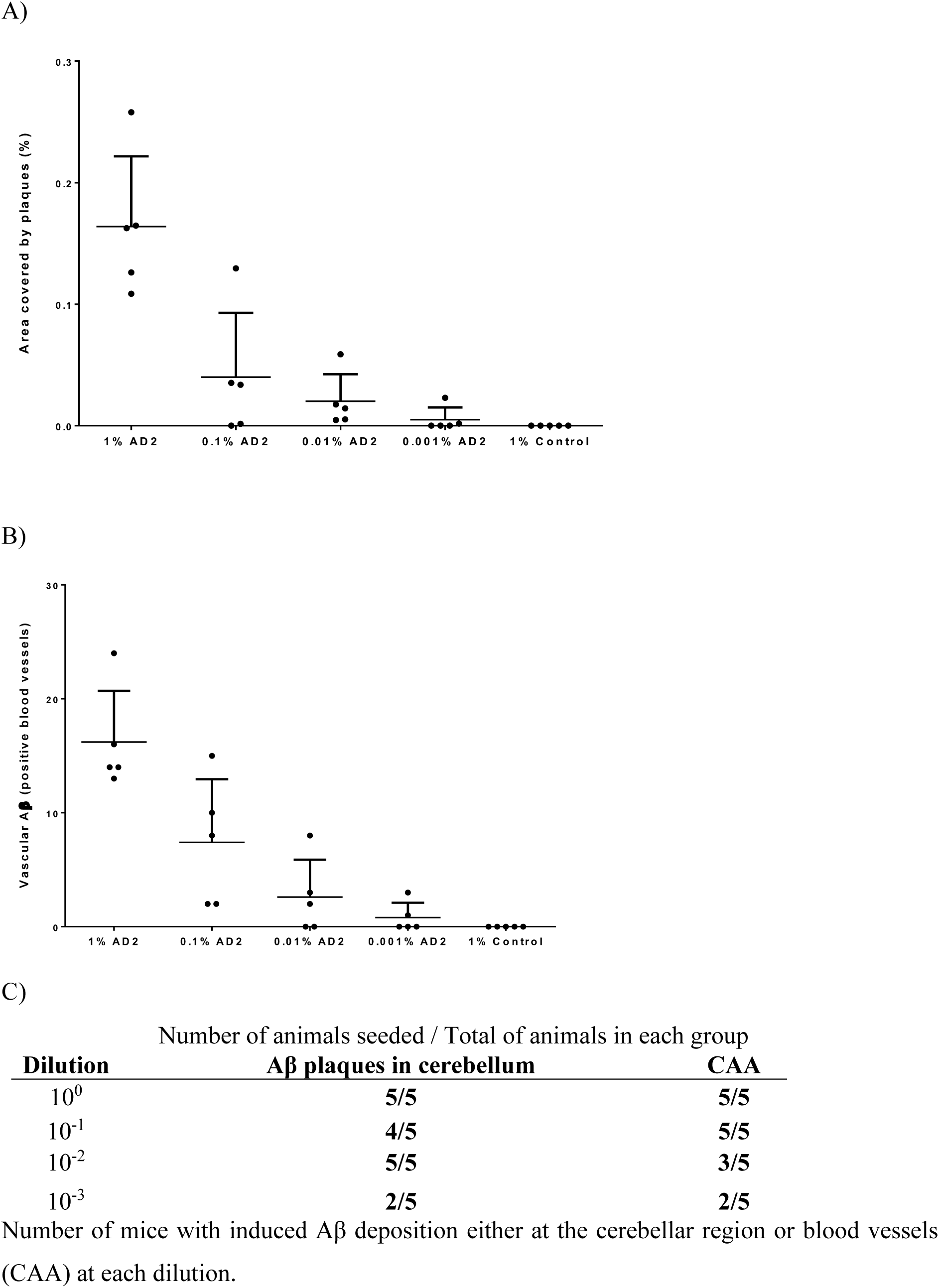
Quantification of cerebellar and meningeal amyloid in NLF mice inoculated with serial dilutions of AD brain homogenate. Four months after inoculation with serial dilutions of control or Alzheimer’s disease (AD2) brain, 5 mice per group were culled and one sagittal section per mouse was stained with biotinylated 82E1. A) Control-inoculated mice were significantly different from AD2 1 % (P<0.0001) regarding the coverage of cerebellar region with Aβ plaques. B) Control-inoculated mice presented highly significant differences (P<0.0001) in CAA count compared to mice inoculated with 1 % AD brain homogenates and with 0.1 % (Control vs 0.1 % P=0.01). One-way ANOVA followed by Dunnett’s multiple comparison test. Mean ± SD is shown. C) Summary of end-point titration.

### Inoculation of NLF mice with AD brain extracts causes tauopathy

Data from natural history studies in Alzheimer’s disease indicate that amyloid deposition precedes and is necessary for the formation of tau tangles, and that disease-associated phosphorylation of tau occurs after amyloid deposition, but before the appearance of tangles (Janelidze, Berron et al. 2021, Luo, Agboola et al. 2022). Similarly, injection of synthetic Aβ into the brains of certain tau transgenic stimulates the accumulation of phosphorylated tau (Gotz, Chen et al. 2001).

We found that even at advanced age NLF mice inoculated with PBS or control brain homogenate show no evidence of AT8 IR, whereas NLF mice inoculated with AD brain show progressive accumulation of AT8 IR in the corpus callosum starting around 120 dpi. Tau pathology was absent in PBS- and control brain-inoculated mice even after the 480 dpi. The three different AD inocula induced formation of AT8-positive aggregates in mice, demonstrating that induction of tau phosphorylation by AD brain homogenate is a generalisable effect. As the AD-inoculated mice aged, levels of AT8-positive tau pathology increased in the corpus callosum, spreading along the ventricles (Figure 7).

**Figure 7.**
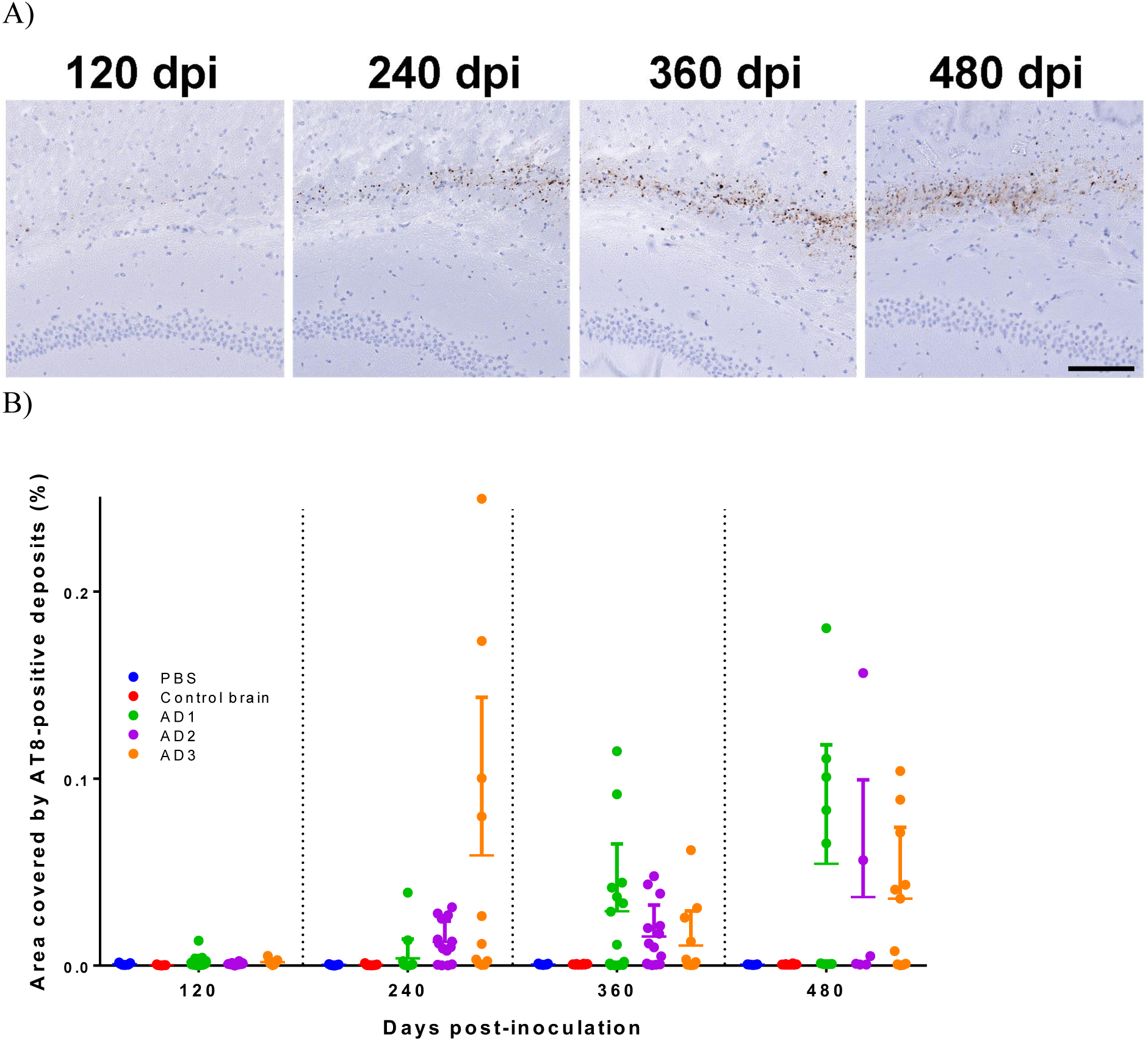
Inoculation of NLF mice with AD brain homogenate induces disease-associated phosphorylation of tau. Sagittal sections stained with anti-phosphorylated tau antibody (AT8) were analysed AT8 depositions were counted. A) Representative images of mice inoculated with AD brain homogenate displaying AT8-positive neuropil thread tau pathology which is not present in PBS- or control brain homogenate-inoculated mice (not shown). AT8-positive pathology was assessed at 120 dpi, 240 dpi, 360 dpi and 480 dpi. B) Quantification of the hippocampus and cerebral cortex area covered in AT8-positive deposits. Mean ± SD is shown. Scale bar: 0.1 mm.

To further investigate why some AD brain inoculated mice are AT8-negative, we investigated whether the levels of Aβ IR were related to accumulation of phosphorylated tau. Direct comparison of AT8 and 82E1 IR in whole brain revealed no simple correlation. However, we found a positive correlation between the area covered by AT8 depositions and the area covered by amyloid plaques when circumscribing our analysis to the hippocampus and cerebral cortex regions (Figure 8). This suggests that tau phosphorylation is accelerated by seeded Aβ pathology in AD-inoculated mice.

**Figure 8.**
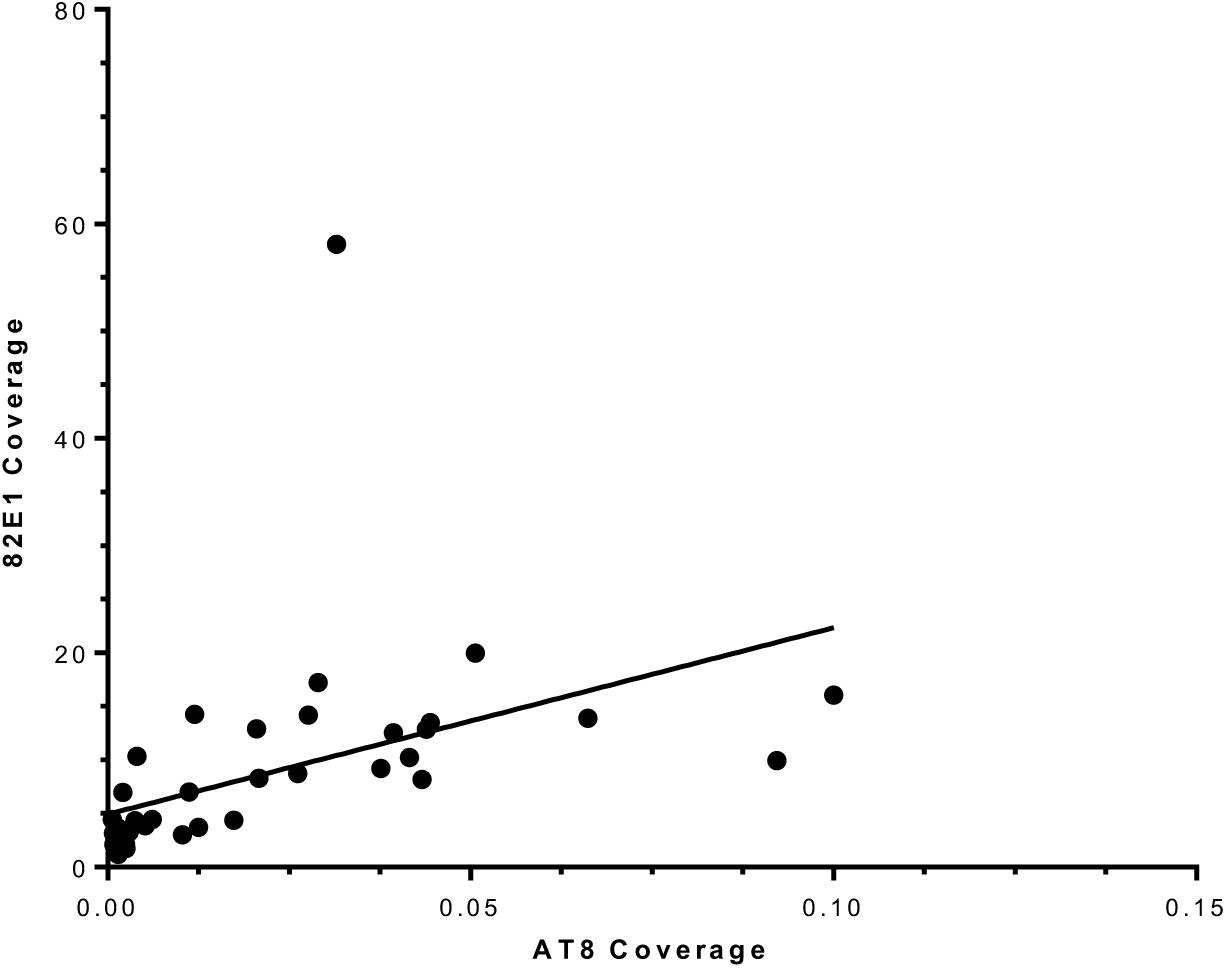
Novel AT8 IR associates with 82E1-positive deposition in NLF mice inoculated with AD brain at 360 dpi. Simple linear regression analysis of the area of the hippocampus and cortex covered by AT8-positive deposits plotted against the area of the hippocampus and cortex covered by 82E1b-positive deposits. A positive correlation was observed in NLF mice inoculated with AD1-3 brain extracts. n=41, Spearman correlation test r=0.81, p<0.0001.

## Discussion

In this study, we have inoculated brain homogenate from three AD patients and one control individual into a knockin AD mouse model expressing Amyloid Precursor Protein (APP) containing the Iberian and Swedish mutations and which produce human Aβ peptides. We performed a detailed longitudinal analysis of Aβ deposition and tau phosphorylation over 480 days post-inoculation, which demonstrated that the NLF knockin mice are a valuable bioassay system to detect and measure Aβ seeds via quantification of amyloid in the cerebellar region, to report vascular Aβ induction and to assess increased Ser202/Thr305 tau phosphorylation and aggregation.

Tau pathology following inoculation with AD brain homogenate has been mainly studied in AD transgenic mouse models with tau pathology, where the injection accelerates Aβ/tau or both pathologies (Robert, Scholl et al. 2021). There are only a few studies using mice expressing murine wild-type tau inoculated with human Aβ+tau seeds and these do not report tau pathology (Kane, Lipinski et al. 2000, Morales, Duran-Aniotz et al. 2012, Ruiz-Riquelme, Lau et al. 2018), except one. Minimal tau hyperphosphorylation in axons of the corpus callosum that had been injured by the injection, following the inoculation with AD brain homogenate in the hippocampus by Walker and colleagues (Walker, Callahan et al. 2002). We show for the first time that inoculating AD brain homogenate in non-transgenic mice accelerates Aβ pathology while murine wild-type tau is phosphorylated and aggregated several months after the Aβ plaques. This is reminiscent of Alzheimer’s disease, in which amyloid plaques and tau tangles co-occur with distinct temporal and topological patterns. Amyloidosis has been suggested as an important driver for the build-up of tau aggregates (Price and Morris 1999) and recently supported by a longitudinal tau PET study, where tau accumulation was only seen in the presence of abnormal amyloid (Jack, Wiste et al. 2018). In agreement with this concept, our study reveals an increase in disease associated tau in regions featuring early seeded amyloid plaques, such as the corpus callosum, in young mice inoculated with AD brain homogenates from 120 dpi onwards. The spread of phosphorylated murine tau progresses slowly, likely due to the species/sequence mismatch between human and murine tau. We would expect more efficient tau seeding upon injection of these brain homogenates in a mouse model expressing human tau given the greater compatibility between the seed and the substrate (Saito, Mihira et al. 2019). From our experiments, it is unclear whether the inoculation of tau seeds present in the human homogenate is sufficient to induce Ser202/Thr305 phosphorylation of murine tau or if the acceleration of Aβ pathology exacerbates human tau seeding and hyperphosphorylation.

During AD, Aβ pathology spreads anterogradely from affected brain regions into areas connected by synaptic pathways (Thal, Rub et al. 2002). In contrast, we show that following intracerebral inoculation with AD brain homogenate, the NLF mice exhibit plaques in the cerebellum as early as 90 dpi. This suggests that the cerebellum is as vulnerable as the cortical region to aggregation of Aβ and injected Aβ seeds can induce a different spatiotemporal pattern of Aβ deposition when compared to their later-onset spontaneous pathology phenotype (Eisele, Bolmont et al. 2009, Eisele, Obermuller et al. 2010). The distribution of Aβ plaques is likely influenced by the initial distribution and processing of these seeds.

Cerebral amyloid angiopathy (CAA) is very commonly found as a co-pathology in people with AD. However, CAA can also present as an independent disease characterised by parenchymal and leptomeningeal vascular Aβ aggregation, causing cerebral haemorrhage and vascular dementia (Biffi and Greenberg 2011). Moreover, iatrogenic CAA (iCAA) has recently been described in individuals with prior exposure to cadaveric human growth hormone or dura mater or who had childhood neurosurgical procedures (Jaunmuktane, Mead et al. 2015, Jaunmuktane, Quaegebeur et al. 2018, Purro, Farrow et al. 2018, Banerjee, Adams et al. 2019, Banerjee, Samra et al. 2022). Inoculation with AD brain homogenate causes a rapid and significant Aβ deposition in meningeal dorsal blood vessels of NLF mice. For two key reasons the NLF mice are an effective reporter system for the induction of vascular Aβ. Firstly, control animals consistently exhibit low numbers of blood vessels containing Aβ deposits throughout their lifespan. Second, the intracerebral injection of Aβ seeds results in a fast deposition in blood vessels, evident as early as 60 dpi. These two factors provide a robust context for the *in vivo* study of Aβ deposition in the vasculature, making this mouse model a helpful tool.

Notably, despite variations in the quality and quantity of total Aβ peptides among the three AD brain homogenates used for inoculation, they induced similar density and distribution of amyloid plaques within comparable timeframes and locations. This suggests that the biochemical quantification of Aβ_38_, Aβ_40_ and Aβ_42_ peptides may not necessarily correlate with seeding potency. Successful seeding may require specific template characteristics akin to conformational selection with prion strains (Collinge and Clarke 2007). Until the full biophysical attributes of an Aβ seed are elucidated (and any co-factor/s), the quantification of Aβ peptides in the inocula should be interpreted cautiously, and cellular bioassays may offer more insightful results (Aoyagi 2019), if an *in vivo* bioassay (titration) is not available.

Notably, only one laboratory previously attempted to titrate Aβ seeds of human origin in mice (Fritschi, Langer et al. 2014). Unfortunately, it is not possible to calculate a SD_50_ from those experiments because the dilutions used did not reach the end-point limit. Collectively, our experiment using AD brain homogenates and others employing AD mouse models brain tissue (Fritschi, Langer et al. 2014, Morales, Bravo-Alegria et al. 2015, Ye, Rasmussen et al. 2017) confirm the titrable nature of Aβ seeds. The ability to assess the minimum amount of biologically active Aβ seeds capable to induce pathology is critical to study the transmissibility of AD. Titration of Aβ seeds present in different biofluids and tissues from AD and related dementias could help to understand the progression of the disease.

Overall, our findings underscore the prion-like propagation of Aβ and tau under physiological conditions and support using the *App^NL-F/NL-F^* mice for further investigations into the mechanisms of Aβ and tau toxicity.

## Acknowledgements

This work was funded by the UK Medical Research Council (MRC); the National Institute of Health Research (NIHR) University College London Hospitals (UCLH)/University College London (UCL) Biomedical Research Centre; the Leonard Wolfson Experimental Neurology Centre and the Foundation for Neurologic Diseases. We thank the Queen Square Brain Bank for Neurological Disorders (supported by the Reta Lila Weston Trust for Medical Research, the Progressive Supranuclear Palsy (Europe) Association and the MRC) at the UCL Institute of Neurology, University College London; and the Oxford Brain Bank (supported by the MRC, the NIHR Oxford Biomedical Research Centre and the Brains for Dementia Research programme, jointly funded by Alzheimer’s Research UK and Alzheimer’s Society) for providing the UK human brain tissue samples. We thank staff of the MRC Prion Unit Biological Services facility for animal inoculation, observation and care and the Unit Histology facility for technical assistance. We thank J. Wadsworth for assistance in selecting and processing tissue samples. Antibody m266 was a gift from P. Seubert and D. Schenk, Elan Pharmaceuticals. Antibodies HJ7.4 and HJ2 were gifts from D Holtzmann. Antibody 1C22 was provided by D Walsh (Harvard). We thank S Mead for helpful discussions. We thank T Saito and T Saido for the NLF mice.

