## Supplementary figures for "Induction and characterisation of Aβ and tau pathology in *App^NL-F/NL-F^* mice following inoculation with Alzheimer’s disease brain homogenate"

Suppl Figure 1:

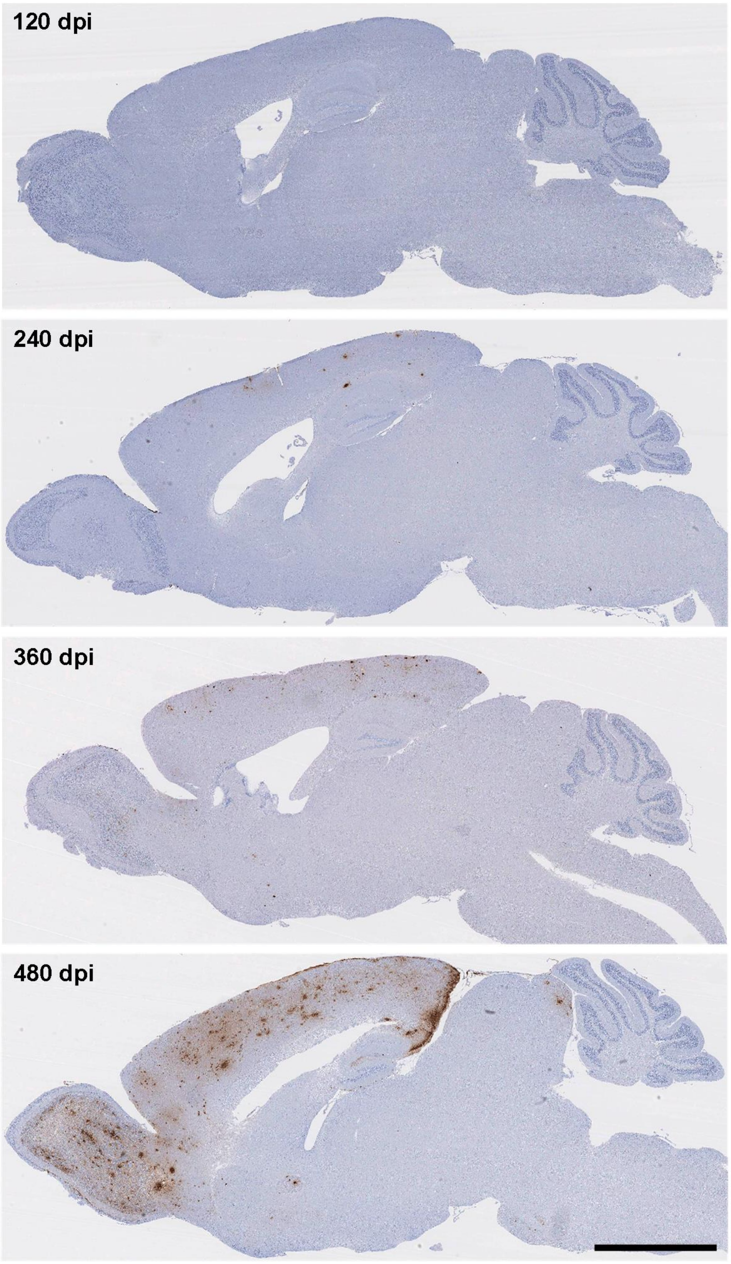


**Progression of endogenous amyloid deposition in PBS-inoculated mouse.** Mice were intracerebrally inoculated with PBS and culled at the times indicated. Aβ deposition was assessed on sagittal sections stained using an anti-Aβ antibody, a biotinylated derivative of the 82E1. Scale bar = 2 mm. dpi = days post-inoculation.

Suppl Figure 2:

A) B) C)

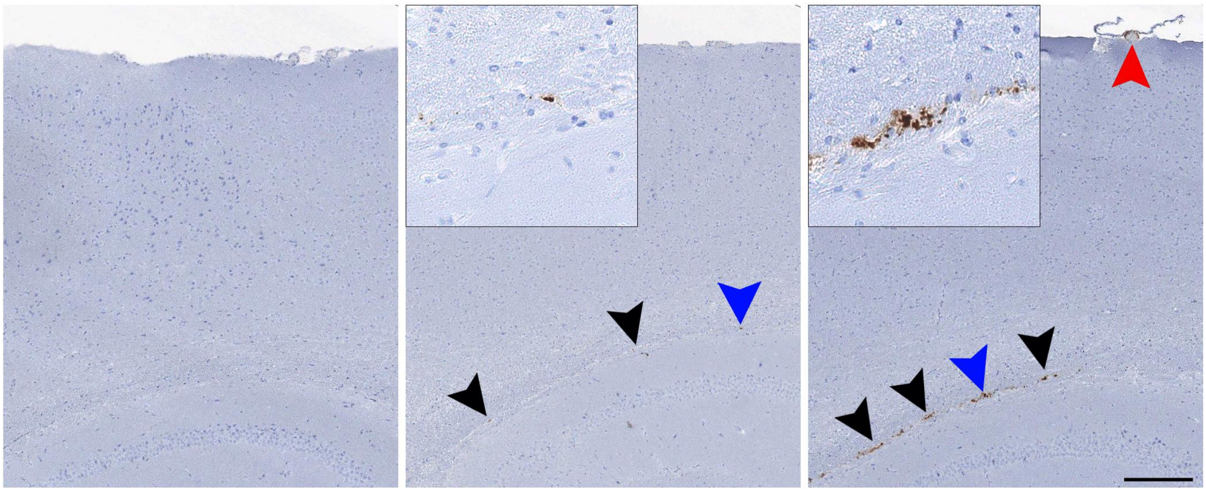

**Progression of amyloid deposition in AD-inoculated mice.** At 2 dpi (A) there was no discernible Aβ. From 30 (B) to 60 (C) dpi few plaques and a couple of dorsal blood vessels with amyloid deposition were already present in the brains of some of the AD1-3 inoculated mice (6 out of 15 mice at 30 dpi and 12 out of 15 at 60 dpi). Red arrow indicates vascular Aβ, black arrows indicate parenchymal Aβ and insets correspond to the areas indicated by blue arrows. Scale bar: 0. 15 mm.

Suppl Figure 3

A)

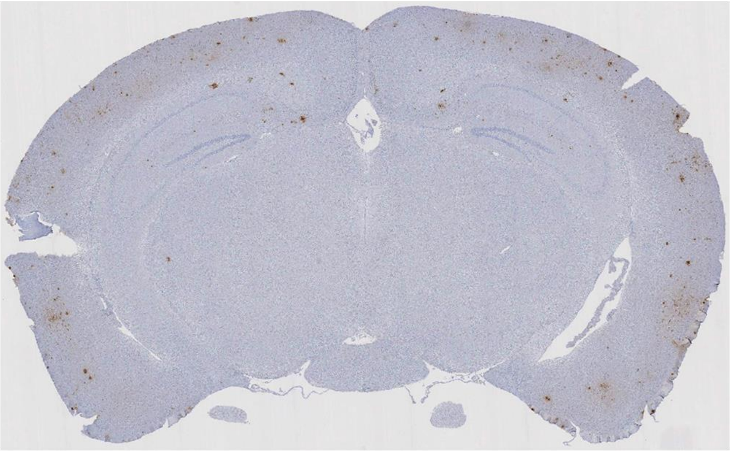

B)

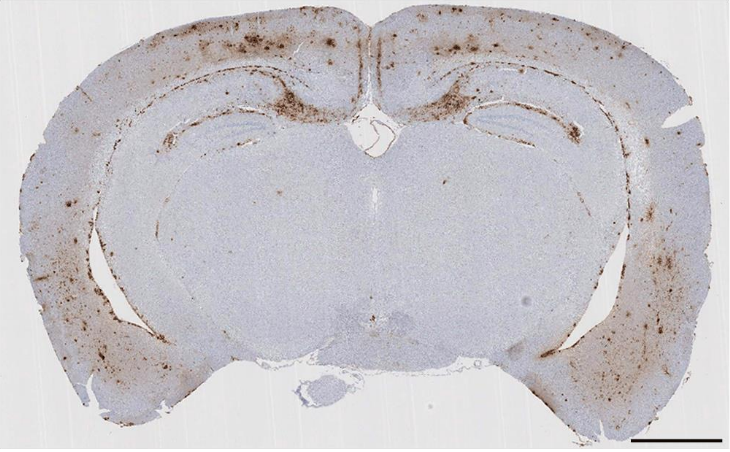

**Comparison of amyloid load on both hemispheres following injection of brain homogenates.** Samples at 120, 240 and 360 dpi were fixed and three coronal cuts were performed to produce four sections from each mouse brain. One slice from each section was stained to quantify the Aβ plaques in each hemisphere. The amyloid load on the ipsilateral side was similar to the contralateral hemisphere in all samples at all time points regardless of the inoculum used. Representative images of one mid-section (bregma -1.7 mm) of Control- and AD3-brain inoculated samples (A and B, respectively) collected at 360 dpi and stained with biotinylated 82E1 antibody. Scale bar: 1 mm.

Suppl Figure 4A, 4B and 4C:

A)

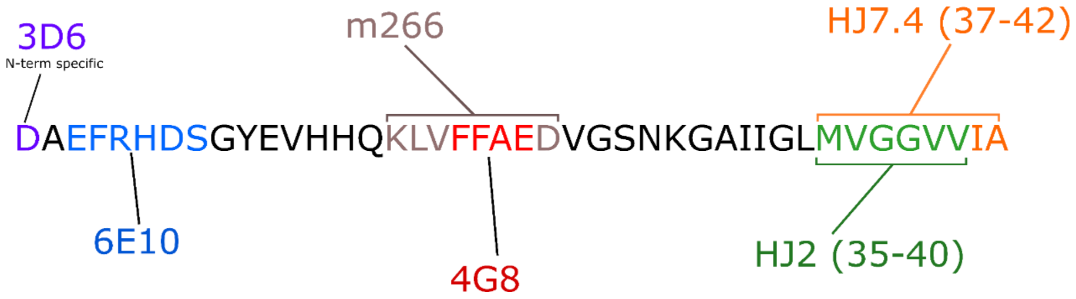


B)

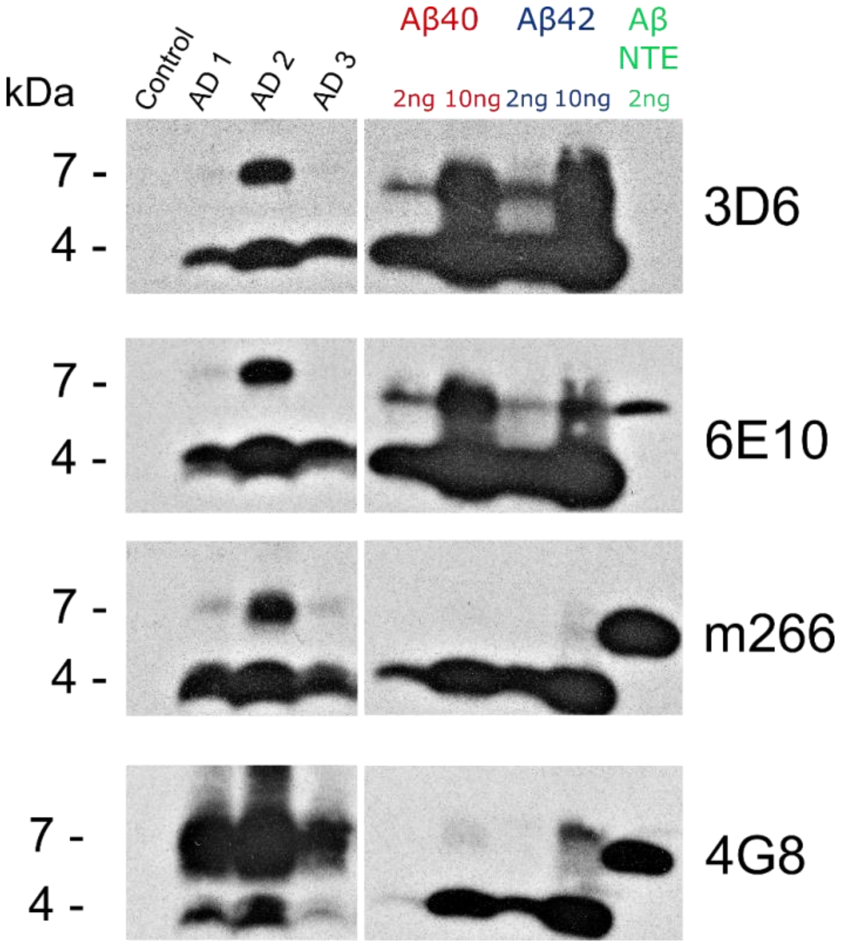

C)

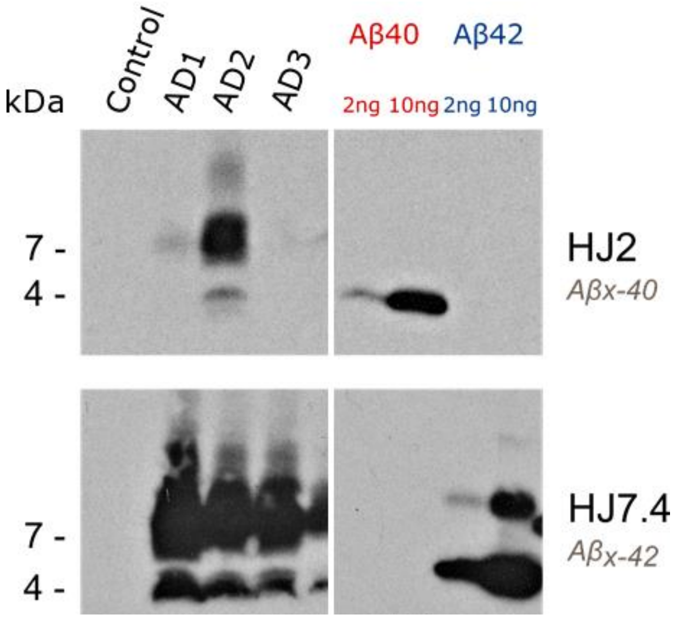


**Western blot analysis of AD brain homogenates using N-terminus, mid-region and C-terminus specific anti-Aβ antibodies.** A) Binding sites for anti-Aβ antibodies: 3D6, HJ7.4 and HJ2 are end-specific to either the N-terminus or C-terminus of specific Aβ peptide variants, meaning they will not bind to full length APP as they bind to the epitope revealed by β- or γ-secretase cleavage. In comparison, 6E10, m266 and 4G8 all bind to epitopes within the Aβ domain that will also be present within full length APP and other APP metabolites and thus will cross-react with these species.

B and C) Human brain homogenates from control and Alzheimer's disease (AD1, AD2, AD3) were thoroughly resuspended and then diluted directly in sample buffer containing 10 % SDS, boiled at 100 $^{o}$C for 10 min, centrifuged at 13,000 rpm for 10 min and the supernatant directly loaded on Tricine gels. 2 ng of NTE $A\beta_{1\text{-}x}$, 2 ng and 10 ng of synthetic $A\beta_{1\text{-}40}$ and $A\beta_{1\text{-}42}$ peptides were loaded on gels to allow comparison across Western blots.

Suppl Figure 5:

A)

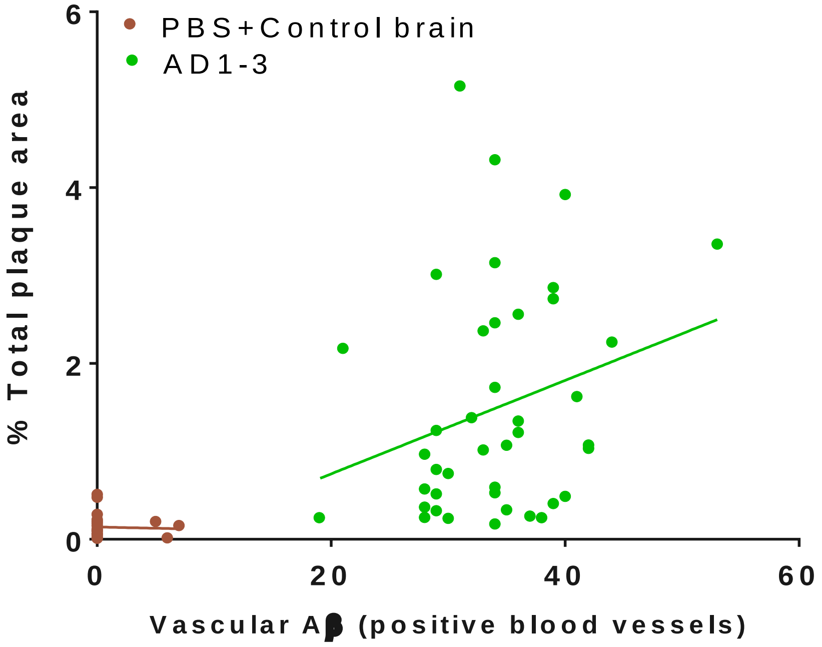


B)

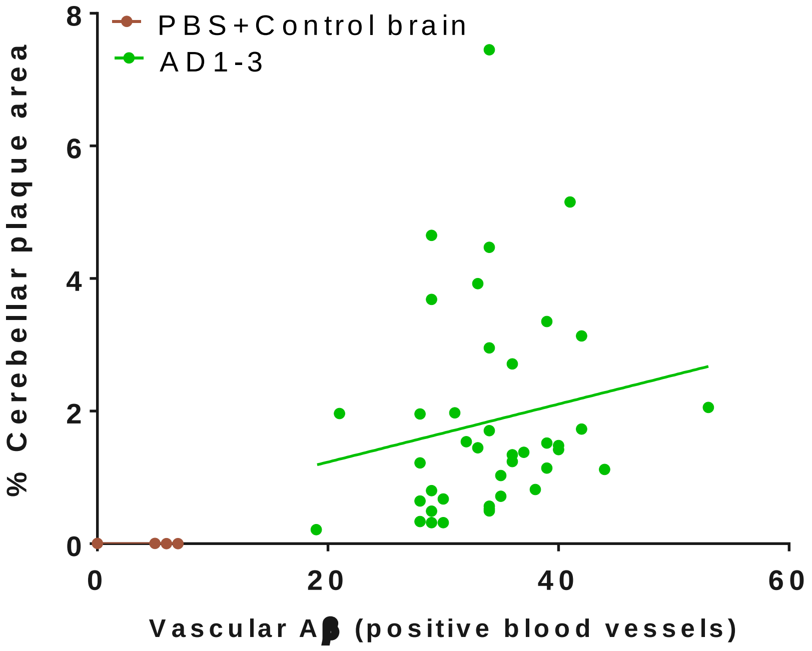


**Correlation analysis of total and cerebellar amyloid plaque burden versus vascular Aβ deposition at 240 dpi.** NLF mice were inoculated with PBS, control brain or AD brain homogenates cases 1-3 and culled at 240 dpi. Sagittal sections were stained with biotinylated 82E1. Dorsal vessels with Aβ immunoreactivity were counted and parenchymal Aβ deposition in the cerebellar region and the total area of the section was quantified. A) Correlation of total plaque area versus vascular Aβ. PBS+Control brain inoculated mice n=25, Pearson correlation test r=0.24, p=0.2. AD1-3 inoculated mice n=41, Pearson correlation test r=0.17, p=0.3. B) Correlation of cerebellar plaque area versus vascular Aβ. PBS+Control brain inoculated mice n=25, Pearson correlation test r=-0.04, p=0.8. AD1-3 inoculated mice n=41, Pearson correlation test r=0.26, p=0.1.
